# The specificity of different-distance connections in human structural connectomes

**DOI:** 10.1101/2022.07.09.499310

**Authors:** Yaqian Yang, Yi Zheng, Yi Zhen, Shaoting Tang, Hongwei Zheng, Zhiming Zheng

**Affiliations:** Institute of Artificial Intelligence, Beihang University, Beijing 100191, China; School of Mathematical Sciences, Beihang University, Beijing 100191, China; Key laboratory of Mathematics, Informatics and Behavioral Semantics (LMIB), Beihang University, Beijing 100191, China; State Key Lab of Software Development Environment (NLSDE), Beihang University, Beijing 100191, China; Beijing Academy of Blockchain and Edge Computing (BABEC), Beijing 100085, China; PengCheng Laboratory, Shenzhen 518055, China; Institute of Medical Artificial Intelligence, Binzhou Medical University, Yantai 264003, China

**Keywords:** structral connectome, specificity, spatial embedding, hierarchy, segregation and integration

## Abstract

Brain structural connectomes underpin complex cognitive processes. To date, abundant organizational features have been distilled by network-based tools, including hubs, modules, and small-worldness. However, these features are often devoid of spatial characteristics which directly shape connection formation. By considering the spatial embedding of brain networks, we reveal the connection specificity, that is, the similarity of similar-distance connections and the dissimilarity of different-distance connections. It is induced by the whole-brain connection length distribution, allowing areas to send and receive diverse signals through different-distance connections. Based on it, areas’ functional repertoires are associated with their connection length profiles, and meanwhile, length dispersion and clustering coefficients can be integrated into a hierarchy whose age-related degeneration may be related to cognitive decline. These results construct a putative bridge between brain spatial, topological, and functional features, expanding our understanding of how different architectures complement and reinforce each other to achieve complicated brain functions.

## 1. Introduction

The brain can be considered as a spatially embedded network composed of interconnected neuronal populations [1]. Interregional connections promote signal transmission and information interaction within the brain network, engendering complicated neuronal coactivation patterns and adaptive cognitive processes [2, 3, 4]. Understanding the architecture of structural connectomes and its functional implication is an ongoing challenge in network neuroscience [5, 6]. Emerging tools from graph theory and network science have been employed to distill essential organizational features, including hubs [7, 8], modularity [9, 10], and small-worldness [11, 12]. These features are believed to support the balance between local, specialized information processing and global, efficient information transmission [13, 14], providing important insight into the organizational basis for advanced cognitive functions. However, network-based approaches often disentangle the brain network from its spatial embedding which directly shapes connection formation.

The spatial embedding of the brain network imposes constraints on axonal outgrowth, drives the expression of significant topological features, and ultimately impinges on brain function [15, 16, 17]. Growth trajectories of most axons are confined to small spheres of space due to the spatial distribution of molecules [18, 19, 20, 21], leading to the probability of brain regions being connected decreasing rapidly with the interregional distance [22, 23]. This dominance of short-range connections reduces the material and metabolic consumption required to construct and maintain connections, facilitating the propagation and preservation of network architecture within surviving species under the pressure of limited time, space, and resources [24]. The physical instantiations of structural connectivity inevitably come with material and metabolic burdens. It has been widely accepted that brain networks are organized under a trade-off between cost constraints and functional demands. Generative models [25, 26] that combine wiring cost minimization and functional complexity requirements reproduced many topological features observed in empirical brain networks, confirming the important roles of spatial embedding and functionality in shaping structural connectomes. Furthermore, the spatial, structural, and functional characteristics of brain networks are implicated with each other. For instance, neurons with similar functions tend to possess similar connectivity profiles [27, 28], and the similarity of areas’ connectivity profiles tend to decrease with interareal distance [29]. Recently, a tight coupling relationship between brain structural and functional connectivity [30] and a macroscale gradient that linked the topographic organization to an increasingly abstract functional spectrum [31] have been revealed. These findings suggest that a holistic understanding of brain organizational mechanisms requires an analysis that simultaneously assesses different aspects of brain intrinsic organization.

Incorporating spatial embedding into network analyses, we showed that the similarity of a brain area’s neighbors decreases with increasing differences in connection lengths. Such similarity of similar-distance connections and dissimilarity of different-distance connections suggest that

This property enables brain areas to receive and deliver unique signals through different-distance connections, which we recapitulated as the connection specificity.

suggesting that different-distance connections may serve to transmit unique and specific signals in interareal communication.

revealed the specificity of different-distance connections, that is, the dissimilarity of different-distance neighbors and the similarity of similar-distance neighbors. Through network randomization techniques, we found this connection specificity was induced by the whole-brain connection length distribution. We then performed individual analyses in two independent datasets to test whether the connection specificity served as a common and fundamental property of brain networks. Since the connection specificity conferred brain areas the capacity to send and receive unique signals through different-distances connections, we assessed the association between connection length profiles and areal functional repertoires as well as the relationship between length dispersion and clustering coefficients. These two issues not only explore how brain spatial, topological, and functional characteristics complement and reinforce each other, but also provide insights into organizational mechanisms underlying diverse information processing. We finally examined the age-related changes in the detected hierarchical structure to explore physiological alterations involved in cognitive decline.

## 2. Results

We explored the spatial characteristics of human structural connectomes and attempted to answer the following questions: How do structural connections with different distances promote and constrain interregional communication? What role does connection length play in shaping areal functional repertoires? How do spatial characteristics harmonize with other characteristics to support diverse information processing?

### 2.1. Spatial influence on the human brain connectivity

We constructed a group-consensus brain connectivity matrix whose elements were colored from blue to red with increasing connection length (Fig. 1A). We observed that the structural network comprises substantial short-distance connections with the admixture of a small number of long-distance connections. We then estimated the wiring length distribution (Fig. 1B) and found that connecting probability exponentially decreased with interregional distance. Finally, we computed the cosine similarity of connectivity profiles of pairwise regions and associated it with interregional distance (Fig. 1C). We found a significantly negative correlation between cosine similarity and interregional distance (*r* = –0.46, *p* < 10^−15^). That is, physically nearby areas tend to have more similar connectivity profiles than physically distant areas. These observations collectively demonstrate the powerful role of interregional distance in shaping brain structural organization.

**Figure 1:**
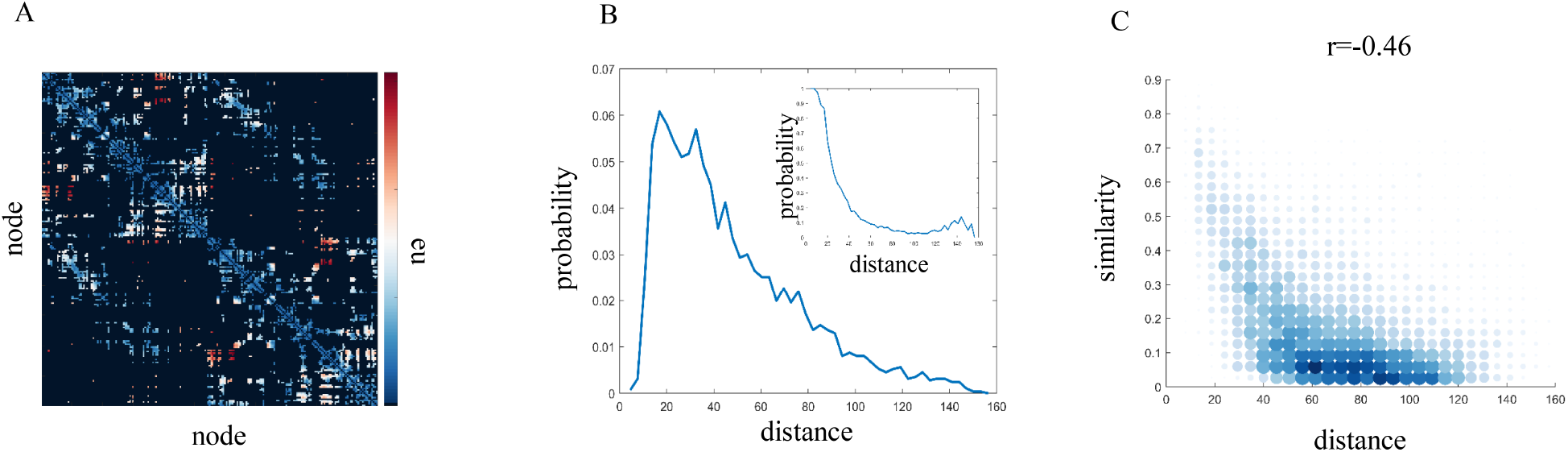
Spatial architecture of the human brain connectome. A. Structural matrix with color corresponding to connection length. Brain structural network contains a large amount of short-range connections with an admixture of long-range connections. B. Wiring probability versus distance. C. Cosine similarity versus distance.

### 2.2. Specificity of structural connectivity

The spatial embedding of the brain network engenders heterogeneity in connection lengths. To explore the role of different-distance connections in interregional communication, we estimated the similarity of brain areas’ different-distance neighbors in terms of their connectivity profiles. The connectivity profile of a brain area shapes its interaction pattern with other areas, determining its functional contribution within the brain network [32, 33]. Areas with similar connectivity profiles tend to possess similar functionality, and thus are expected to contribute similar signals in interregional communication [34, 35, 36, 37, 38].

Specifically, we computed the cosine similarity of connectivity profiles of any two neighbors 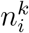 and 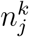 of a given area k, and then partitioned these neighbors into different bins according to their connection lengths. We repeated this process for each brain area and applied the same partition to all areas’ neighbors. To assess the relatedness of inputs and outputs provided by different-distance connections, we computed the average similarity of connectivity profiles of neighbor pairs, for which one neighbor is from bin A and the other from bin B: 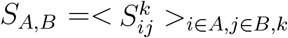. The resulting similarity matrix measures the similar degree to which brain areas receive or deliver inputs and outputs along different-distances connections. Furthermore, we performed a z-transformation of the similarity matrix to have a clearer and more intuitive vision. The processing pipeline is illustrated in Fig. 2A.

**Figure 2:**
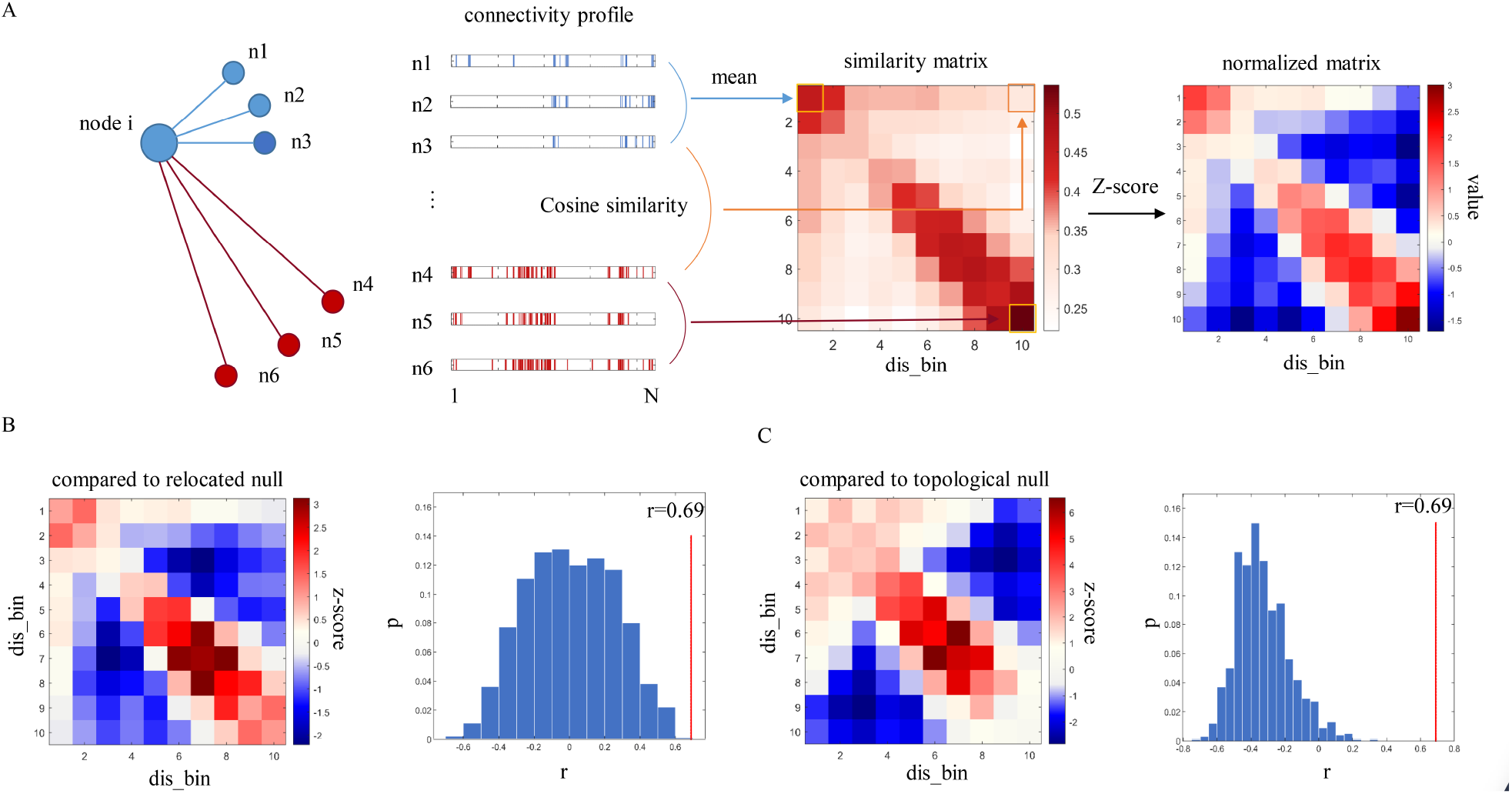
Specificity of different-distance connections. A. Method pipeline for assessing the similarity of neighbors connected by different-distance connections. For any given node i, its neighbors are classified into different distance bins. The connectivity profiles of different-range neighbors are compared with one another using cosine similarity. We repeat this process for each individual nodes and obtain a batch of pairwise similarity measures. Note: the same distance partition is applied uniformly to all network nodes. The similarity matrix is composed of the mean similarity over pairs within or between different distance bins. The z-transform of the similarity matrix is evaluated to have a clearer and more intuitive vision. B. Comparison between empirical values with a null distribution generated by randomly permuting areas’ spatial locations while preserving network topology. C. Comparison between empirical values with a null distribution generated by randomly rewiring network edges while preserving degree sequence.

We observed that positive z-scores are concentrated along the diagonal of the normalized similarity matrix, that is, the similarity of connections with similar lengths is higher than the global level. From brain area k’s perspective, neighbors linked by similar-distance connections have similar connectivity profiles, therefore exhibiting similar functional repertoires and providing a set of redundant input and output signals. This structural degeneracy confers brain areas the capacity to deliver and receive similar signals in the event that a small number of connections are damaged, potentially promoting the system’s resilience to perturbations. In parallel, we found standardized similarity scores between different bins are negative and tend to decrease with increasing differences between connection lengths. This observation indicates that different-distance connections lead to neighbors whose connectivity profiles are more dissimilar than those linked by similar-distance connections, therefore enhancing the diversity of areal inputs and outputs. This dissimilarity of different-distance connections may be an organizational substrate that allows diverse signals to be transformed and integrated in specific brain areas, potentially supporting complex cognitive functions and adaptive human behaviors.

The degeneracy of similar-distance connections and the diversity of different-distance connections allow us to ascribe functional significance to connection distance. For simplicity, we recapitulated these two attributes as the connection specificity. To quantify the extent to which different-distance connections link to dissimilar neighbors and transmit different signals, we measured the specificity degree r as the negative correlation coefficient between similarity scores and connection length differences. Larger values indicate the increased uniqueness of signals transmitted by different-distance connections. To test whether the connection specificity is a meaningful architecture of brain networks, we compared the normalized similarity matrix and r-value observed in empirical data against those obtained from two commonly used null models. In the first null model, we preserved the network topology but permuted areas’ spatial locations. As shown in Fig.2B, the similarity scores are positive for similar-distance connections but negative for different-distance connections, More quantitatively, we observed that the empirical r-value is significantly larger than the null distribution generated by spatial permutation (*p* < 10^−3^). These results indicate that the specificity of different-distance connections is a nontrivial organizational property of the empirical brain network. In the second null model, we compared the empirical values against a null distribution generated by randomly rewiring network edges while preserving the degree sequence (Fig. 2C). We observed that the similarity scores are significantly negatively correlated with the differences between connection lengths (*p* < 10^−3^), confirming that the specificity of different-distance connections is a meaningful organization property that cannot be simply explained by the degree sequence.

### 2.3. Contribution of wiring length distribution

Network randomization techniques have been considered as powerful tools in brain network analysis. Random network null models that retain the number of nodes, number of connections, and degree distribution sever as important benchmarks in identifying meaningful features potentially relevant to biology. Extensions of this approach, via null-hypothesis or surrogate comparison, aid researchers to test whether observed statistics are induced by or at least affected by the particular network property. For instance, one can assess the functional role of brain long-range connections by comparing random surrogates that preserve long-distance connections with those that do not [39]. Here, we focused on the specificity of structural connections and constructed four different types of surrogate networks: (i) random surrogates(R) that disrupt all network properties except the number of nodes and connections; (ii) degree-preserving surrogates(D) that restore the degree sequence of the empirical network; (iii) geometry-preserving surrogates(G) that restore the wiring length distribution of empirical network; (iv) degree-geometry surrogates(DG) that simultaneously restore original degree sequence and wiring length distribution. We capitalized on these four distinct null models to investigate whether the specificity of different-distance connections can be attributed to fundamental network properties conserved in surrogates.

As shown in Fig. 3, spatially constrained surrogates (G and DG) exhibited the architecture of connection specificity while random(R) and degree-preserving surrogates(D) do not. The rvalues that measure the specificity degree were comparable in the empirical network and spatial surrogates (G and DG), but significantly larger than those in random and degree-preserving surrogates (*r_E_* = 0.690, *r_DG_* = 0.782 ± 0.015, *r_G_* = 0.762 ± 0.017, *r_D_* = 0.328 ± 0.156, *r_R_* = 0.006 ± 0.241). These observations indicate that the connection specificity in the structural connectome may be predominately attributed to its wiring length distribution. In addition, considering that all spatially constrained randomized networks (E, DG, and G) exhibit high r-values, we speculated that the distribution of wiring length appeared to be sufficient for the expression of connection specificity, irrespectively of the particular arrangement of structural connections.

**Figure 3:**
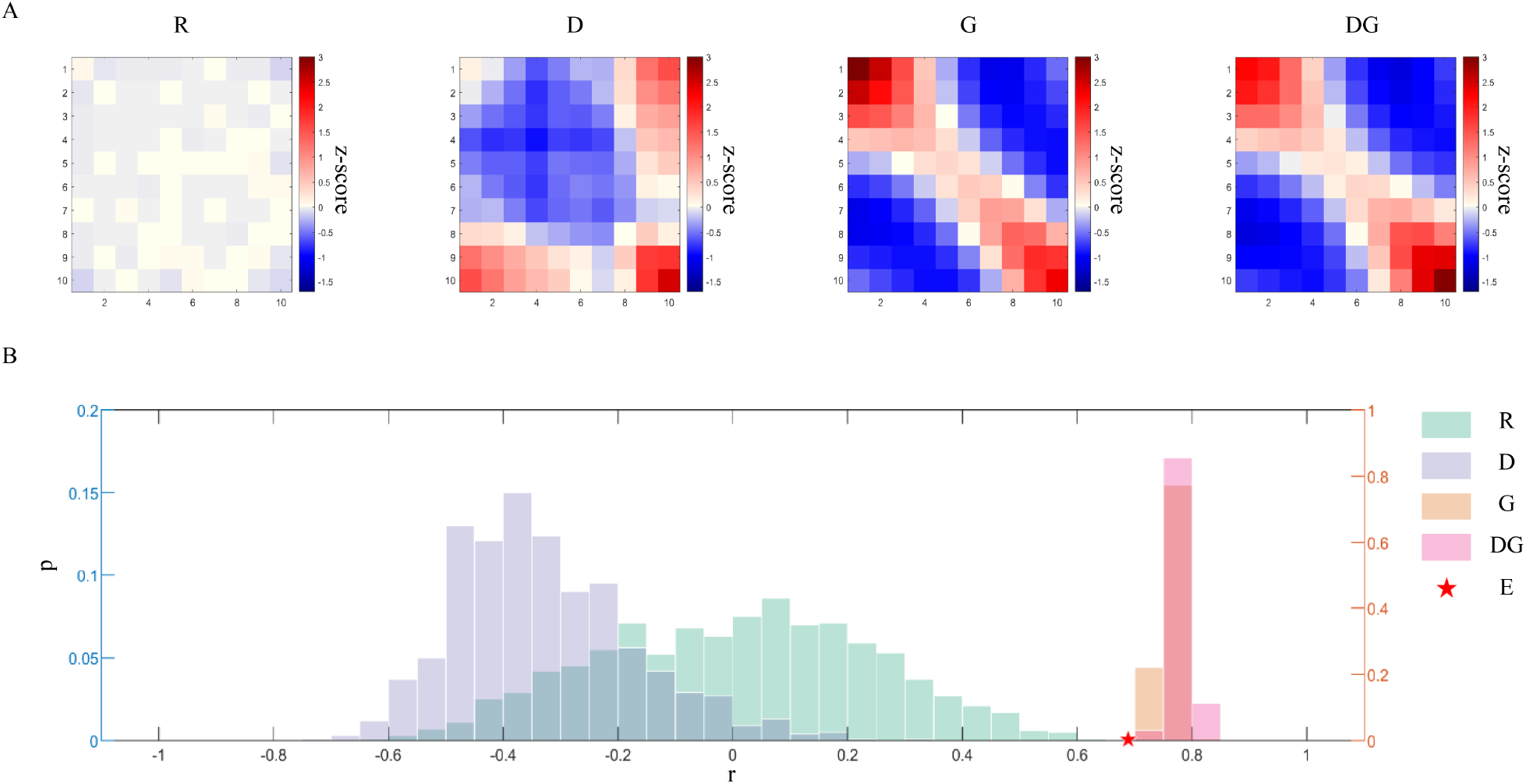
Surrogate-comparison analysis. A. The averages of the similarity z-score matrices of four surrogate groups: (i) random surrogates(R) that disrupt all network properties except the number of nodes and connections; (ii) degree-preserving surrogates(D) that restore the degree sequence of empirical network; (iii) geometry-preserving surrogates(G) that restore the wiring length distribution of empirical network; (iv) degreegeometry surrogates(DG) that simultaneously restore original degree sequence and wiring length distribution. B. The specificity degree r of empirical network(E) and the r distributions of four surrogate groups.

To explore the effect of wiring length distribution on the expression of connection specificity, we introduced degree-preserving perturbations progressively (from 0 to 100%) to the empirical connectome. In the process of partial randomization, the desired fraction of edges was selected for rewiring, generating an ensemble of 100,000 surrogate realizations whose wiring length distributions are interpolated between original and fully randomized networks. The wiring length distribution’s variation degree in each surrogate network was estimated as the Kullback-Liebler (KL) divergence between distributions in the surrogate and the empirical network. As shown in Fig. 4, we found the degree of connection specificity monotonically decreased with increasing K-L divergence. However, the connectivity similarity or overlapping proportion between the surrogate and the empirical network also decayed in a manner coordinated with these two measures, leading to an alternative possibility that the gradual disappearance of connection specificity might be the consequence of incremental randomization. To rule out this possibility, we further applied geometry-preserving randomization to each surrogate network. We found that the relationship between the specificity degree and K-L divergence was almost unchanged despite the substantial decrease in connectivity similarity between the empirical network and these new surrogates (similarity=0.245±0.060). Altogether, these results suggest that the complex dependence of connecting probability on interregional distance entails the expression of connection specificity in physically embedded brain networks.

**Figure 4:**
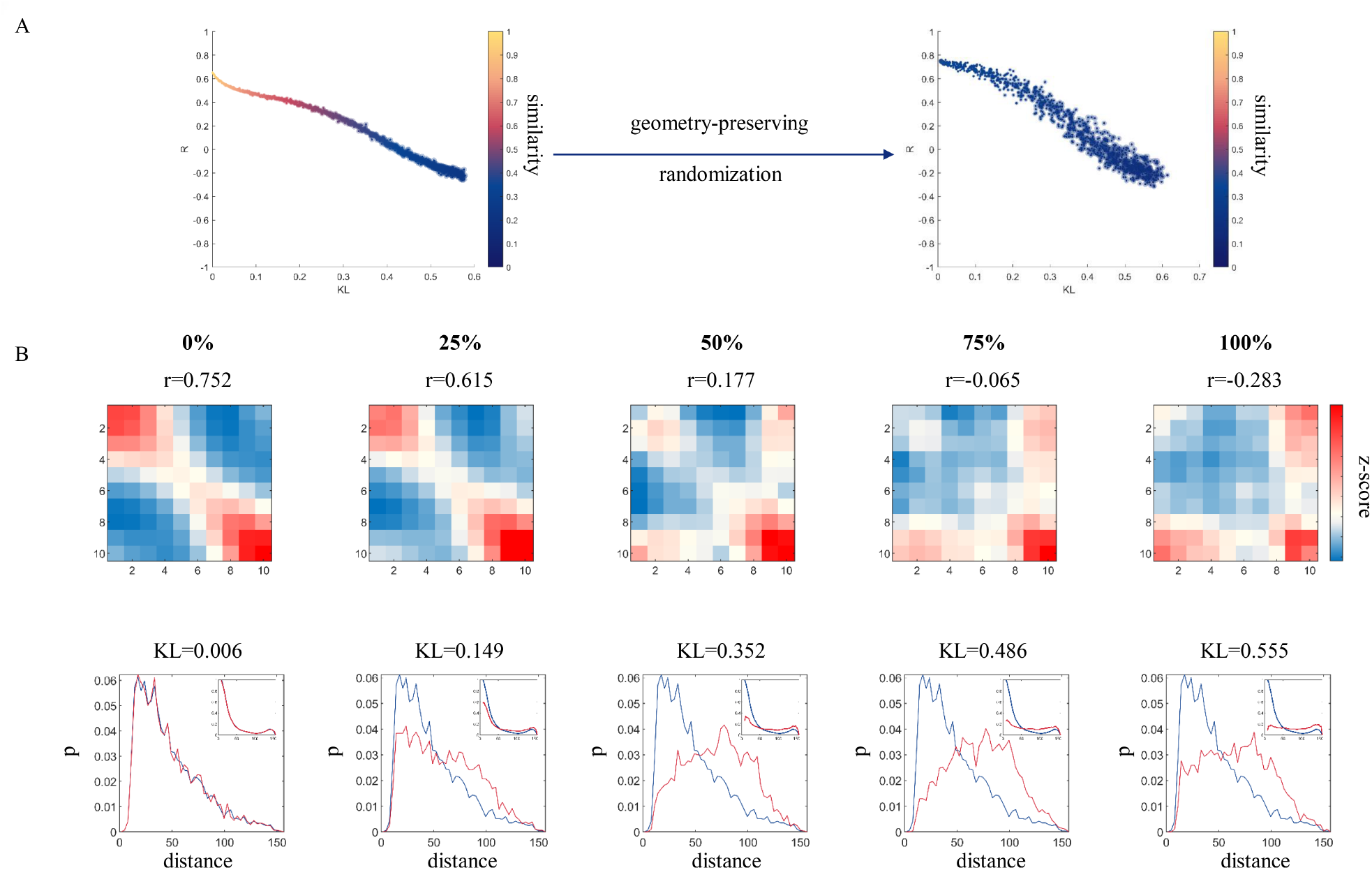
Specificity degree as a function of the wiring length distribution. A. Evolution of connectivity specificity degree r for incremental Kullback-Leibler divergence, with colored points showing the connectivity similarity between different randomized surrogates and the empirical network. B. Standardized similarity matrices(top panel) and corresponding wiring length distributions(bottom panel) of exemplar surrogates that are interpolated between the original and fully randomized networks.

### 2.4. Consistency and variability across individuals

Despite substantial variability in individual structural connectomes, salient organizational features that play crucial roles in supporting brain function are expected to be consistently conserved among healthy individuals. Therefore, we expected the connection specificity to be replicated and conserved across individual participants. Furthermore, since connection specificity in brain networks naturally arises from the spatial constraints irrespectively of the particular arrangement of structural connections, we speculated that wiring length distributions were strictly consistent across individuals while the instantiation of physical connections might exhibit moderate inter-subject variability.

To test the above hypothesis, we estimated the specificity degree of each individual brain network. We also computed the connectivity similarity as well as K-L divergence between each individual network and the group-consensus network. When comparing these measures with a landscape generated by incremental randomization, we found that all individual networks were located within a narrow top-left window in r-KL space (Fig. 5). In other words, all Individual brain networks consistently exhibit a high level of connectivity specificity (*r* = 0.584 ± 0.074) and a low level of variability in wiring length distribution (*KL* = 0.023 ± 0.013). We further found that connectivity similarity of individual brain networks was moderately lower than that of surrogate realizations within this narrow window, indicating a looser constraint of connectivity arrangement relative to the wiring length distribution. Since all individuals are healthy participants, these observations suggest that connection specificity is a common organizational property of human connectomes while moderate connectivity variability would not interfere with normal brain function.

**Figure 5:**
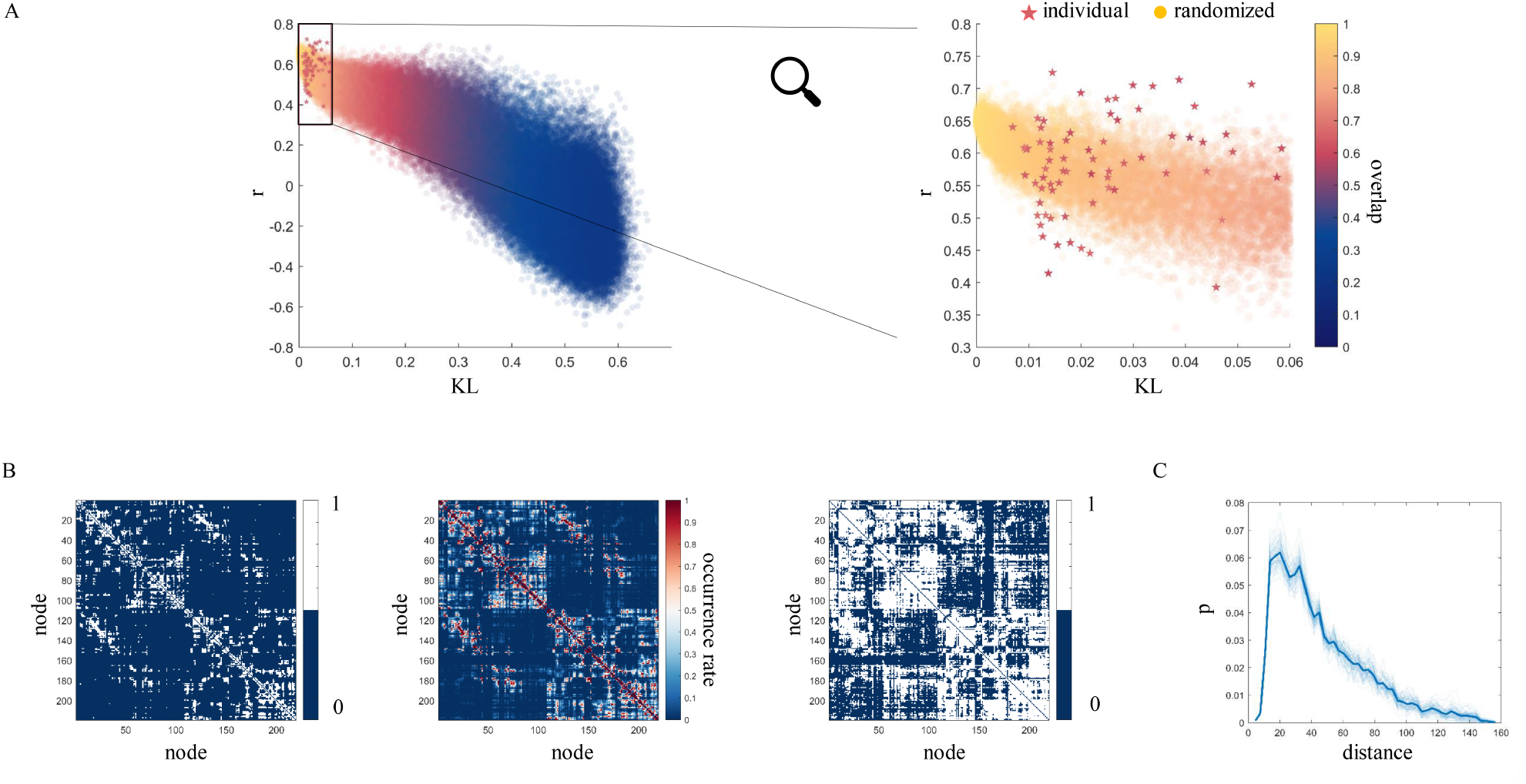
individual analysis. A. Comparison between individual networks (pentagons) and those generated from incremental randomization (circles). B. Structural matrices with edges that consistently appear among all individuals (left panel), with edges weighted by occurrence rates (middle panel), and with edges that appear in at least one participant (right panel). All matrices are derived from 70 healthy participants. C. The consistency of length distributions. The thick line indicates the group-consensus length distribution while transparent lines represent individual length distributions.

### 2.5. Relevance to functional repertoires

Structural connections with different distances transmit unique information in interregional communication, driving the diversity of brain areas’ functionality. It is intuitively possible that connection length profiles mirror the functional repertoires of brain areas. To test the relationship between connection length profiles and areal functionality, we clustered brain areas according to their length distributions using the k-mean clustering method and evaluated the normalized mutual information (NMI) between areas’ clustering assignments and their membership in seven resting-state networks proposed by Yeo et al [40] (Fig. 6A). In concert with prior work [41], we found the NMI values were significantly higher than those generated by randomly clustering assignments (*p* < 10^−3^), implying that areas’ length profiles are informative indicators of their functional roles. The average length distributions of seven functional systems were plotted in Fig. 6B.

**Figure 6:**
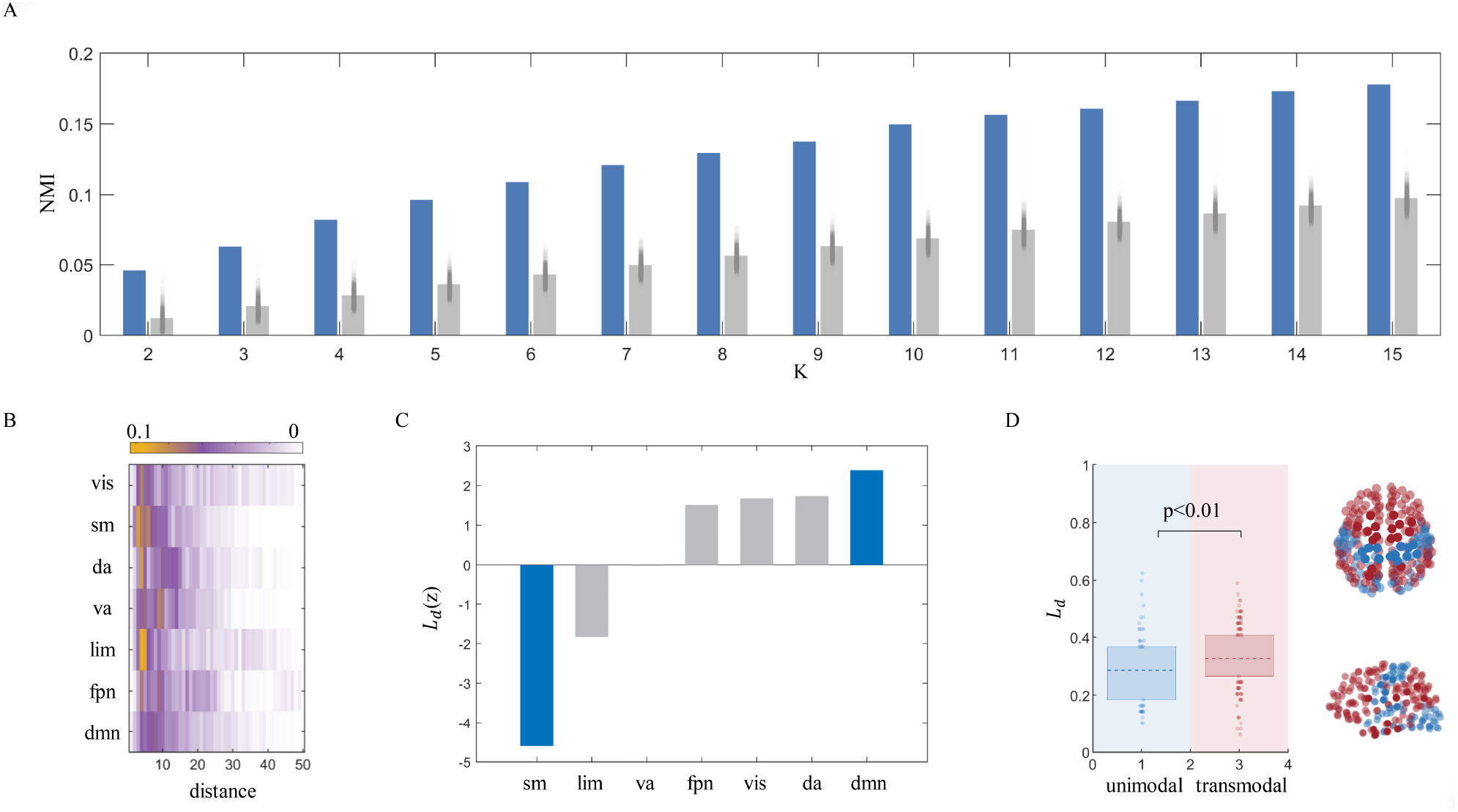
Relationship between connection length distributions and functional repertoires. A. Normalized mutual information (NMI) between areas’ clustering assignments and their membership in functional systems. B. The average length distributions for seven resting-state networks: somatomotor(sm), limbic(lim), ventral attention(va), frontoparietal(fp), visual(vis), dorsal attention(da), default mode(dmn). C. Standardized length dispersion of seven resting-state networks. The network-specific *L_d_* is expressed as a z score relative to a spatial permutation null distribution. Statistically significant networks are shown in blue while nonsignificant networks are shown in gray. D. Comparison between unimodal and transmodal areas. Unimodal areas: areas in visual and somatomotor networks. Transmodal areas: all other areas

More quantitatively, we measured the length dispersion *L_d_* of each brain area (see Methods). The *L_d_* is bounded in [0,1] with the larger value indicating the more uniform length distribution. Aggregating *L_d_* by functional systems, we found that the somatomotor network exhibited significantly lower *L_d_* than expected by chance and that the default mode network exhibited significantly higher *L_d_* than expected by chance (10,000 permutation tests; *p* < 0.05) (Fig. 6C). We further divided seven resting-state networks into unimodal and transmodal groups to assess whether regions associated with functional specialization and functional integration can be distinguished from each other in connection length dispersion. We found that transmodal regions had significantly higher *L_d_* relative to unimodal regions (*p* < 0.01) (Fig. 6D), suggesting that transmodal areas contained more uniform connection length distributions than unimodal regions.

Collectively, these findings demonstrate that brain areas’ length profiles are related to their functionality, with low length dispersion supporting functional specialization and high length dispersion corresponding to functional integration. This observation provides spatial mechanisms of diverse information processing, advancing our understanding of how the structural substrate gives rise to rich functional repertoires.

### 2.6. Hierarchical architecture

The length dispersion *L_d_* varied considerably across the brain areas, ranging from *L_d_* = 0.061 to *L_d_* = 0.624 (Fig. 7A). Some regions have narrow and sharply peaked connection length distributions while others have broad length distributions. Based on the specificity of different-distance connections, we hypothesized that the variability of areas’ length dispersion might be implicated with a segregation-integration hierarchy in the brain network. That is, areas with low length dispersion may be linked to functionally coherent and densely coupled neighbors, thus exhibiting high clustering and participating in segregated information processing. In contrast, brain areas with high length dispersion would be linked to spatially distributed and sparsely interconnected neighbors, thus exhibiting low clustering and serving as the confluence of diverse information.

**Figure 7:**
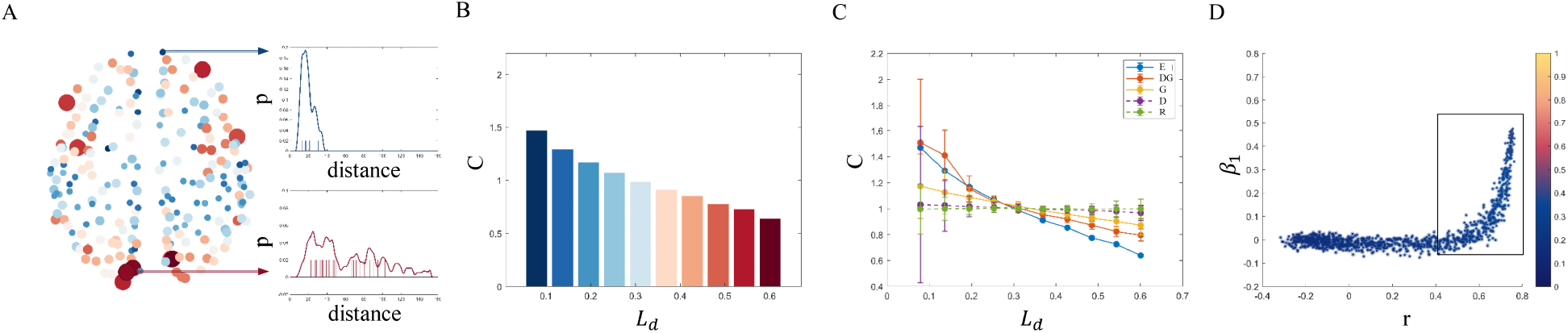
The hierarchical structure. A. Regional connection length dispersion. The orange nodes exhibit large length dispersion while the blue nodes exhibit low dispersion. B. The relationship between length dispersion and clustering coefficients. C. the clustering-dispersion relationship in empirical data vs. four null models: random surrogates(R), degree-preserving surrogates(D), geometry-preserving surrogates(G), and degree-geometry surrogates (DG). D. The effect of connection specificity on network hierarchy.

To test this hypothesis, we characterized the relationship between the connection length dispersion *L_d_* and clustering coefficient C in the brain network. Building on the intuition that nodes contacted with multiple distinct information sources should occupy a central position in the integration, we measured the hierarchical structure as the tendency of nodes with broad length distributions (high *L_d_*) to connect separated parts of the network (low C). Specifically, we divided brain regions into different tiers based on their length dispersion and then computed the average normalized clustering coefficients of areas within each tier. We observed a monotonous decline in the mean clustering coefficient with increasing length dispersion, indicating a well-expressed hierarchical organization of the brain network (Fig. 7B). We further compared the clustering-dispersion relationship in four null models: random surrogates(R), degree-preserving surrogates(D), geometry-preserving surrogates(G), and degree-geometry surrogates (DG). We found that the hierarchical structure emerged in spatially constrained networks (G DG and E) but not in degree-preserving(D) and random surrogates(R) (Fig. 7C), indicating that the hierarchical structure was a nontrivial property induced by brain spatial characteristics.

Since the alignment between length dispersion and the functional integration was derived from the specificity of different-distance connections, we attempted to investigate the effect of connection specificity on network hierarchical structure. We generated 100, 000 degree-preserving random networks and assessed their specificity degree and hierarchy index (see Methods). Interestingly, we found that the hierarchy index *β*_1_ fluctuated up and down around zero until the specificity degree r increased to a relatively large value (approximately 0.4), and then, *β_1_* increased with increasing r (Fig. 7D). This observation suggests that connection specificity may be a prerequisite for hierarchical structure and that the increment in specificity degree could induce a more discriminative hierarchical structure.

Collectively, these findings demonstrate a hierarchical organization of the brain structural network, associating regional heterogeneity in connection length dispersion with a segregation-integration gradient measured by clustering coefficients. Areas with broad connection length distributions tended to have lower clustering coefficients than those with narrow length distributions, and this discrepancy was enhanced as the specificity degree up. This observation provides a novel bridge between spatial characteristics and topological features of the brain network, highlighting the significant role of structural connection specificity in supporting specialized-integrative information processing.

### 2.7. Age-related degeneration of the hierarchy

In the previous section, we detected a hierarchical structure that may serve for the segregated-integrative information processing. We speculated that aging and associated cognitive deficits might be linked to the degeneration of this hierarchy. In this section, we employed the Nathan Kline Institute (NKI)/Rockland Sample public dataset, which included 196 healthy subjects from 4 to 85 years old, to test whether brain hierarchical architecture exhibit age-related alternations in human aging process.

Since the hierarchy of the brain network emerges from the connection specificity, we first assessed the specificity degree of each healthy participant. We found that all individuals had a significantly higher level of connection specificity than degree-preserving surrogates (t-tests, FDR corrected, *p* < 10^−15^) (SI Appendix, Fig. .1) and that the specificity degree was weakly associated with age (r=0.150, p=0.036) (Fig. 8A). This result not only provided a prerequisite for the emergence of hierarchical structure but also confirmed that connection specificity was a common and fundamental property of healthy brain structural networks during various developmental and aging stages.

**Figure 8:**
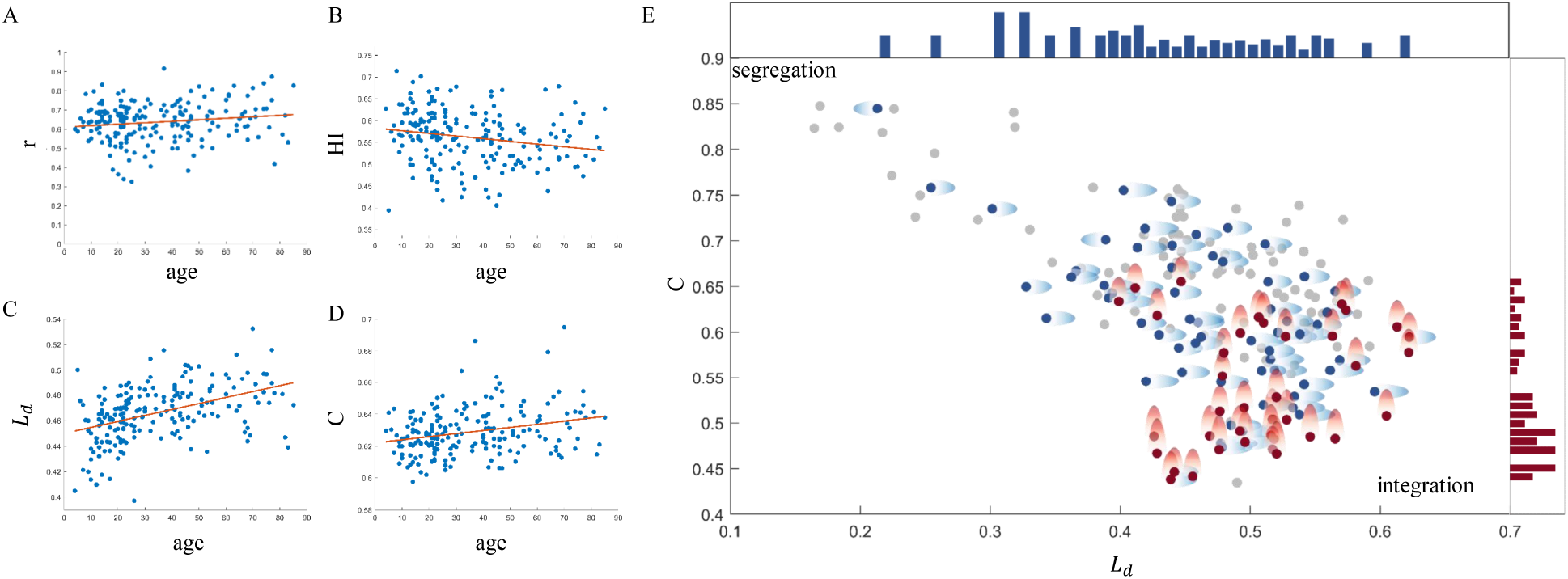
Age-related alteration in the network hierarchy. A. The specificity degree was weakly associated with age. B. The hierarchy index was significantly negatively correlated with age. C&D. Both global length dispersion and global clustering coefficient were significantly positively correlated with age. E. The age effect on length dispersion and clustering coefficients in individual brain regions.

Next, we attempted to investigate how aging affects the hierarchical structure of the brain network. Specifically, we computed the hierarchical index of each individual network and correlated them with participants’ ages. We found a significant negative correlation between the hierarchy index and the age (r=-0.195, p=0.006) (Fig. 8B), indicating the degeneration of hierarchical structure in the aging process. We further plotted the lifespan trajectories of network-average length dispersion and clustering coefficient (Fig. 8C&D). We found that both length dispersion and clustering coefficient were significantly positively correlated with the age (*r_L_* = 0.432, *p_L_* < 10^−9^; *r_C_* = 0.267, *p_C_* < 10^−3^).

Finally, to study how age-related changes in length dispersion and clustering coefficient combined to generate hierarchy degeneration, we analyzed the age effect on these two metrics at the level of individual brain regions. We observed 67 nodes exhibited significant age-related changes in length dispersion and 38 nodes exhibited significant age-related changes in clustering (*p* < 0.05, FDR corrected). Remarkably, these two groups overlapped slightly (with only 8 nodes in common), indicating regional heterogeneity of age effects on these two network attributes. This observation also suggested that although length dispersion and clustering coefficient were strongly associated with each other, they represented different aspects of brain structural organization. When mapping network nodes back to the hierarchical structure (Fig. 8E), we found that nodes whose length dispersion increased significantly with age are broadly distributed across different hierarchical tiers. The increment in length dispersion enhanced the diversity of areal inputs and outputs, potentially promoting areas’ capacity for information integration. However, this alteration might also introduce noise interference to specialized information processing. Besides, we found nodes with significant positive age-related changes in clustering coefficients were concentrated in the integration space (low C and high *L_p_*). These nodes, which were configured to communicate with separated network parts and to integrate diverse signals, were gradually embedded in more homogeneous neighborhoods due to their increasingly interconnected neighbors, thus potentially exhibiting the weakened capacity for intermodular communication and information integration. Collectively, these results demonstrate an age-related degeneration of the segregation-integration hierarchy, providing insights into physiological alterations concerning cognitive decline during human aging process.

## 3. Discussion

Brain function is naturally shaped by the physical architecture; however, it is still challenging to understand how structural connections are organized to support sophisticated cognitive processes. In this work, we adhered to the analysis that simultaneously assesses topological and spatial characteristics of the brain structural network. We showed that, for a given brain area, the connectivity profiles of neighbors linked by similar-distance connections exhibited higher similarity than those linked by different-distance connections, which we recapitulated as the connection specificity. This connection specificity provided an underlying mechanism of interregional communication: brain areas receive and deliver unique signals through different-distance connections. Through network randomization techniques, we further demonstrated that the connection specificity was shaped by the whole-brain wiring length distribution, irrespectively of the particular arrangement of structural connections. In other words, the connection specificity is induced by the spatial architecture of the brain structural network.

If the connection specificity is an essential organizational feature underpinning brain function, it can be expected to be conserved in different healthy individuals. Furthermore, for the sake of the emergence of connection specificity, the wiring length distribution should be strictly consistent across individuals while the instantiation of structural connections could exhibit moderate inter-subject variability. Notably, these speculations are concordant with the findings in individual analysis. All individuals’ brain networks consistently exhibit a high level of connectivity specificity and a low level of variability in wiring length distributions. When contextualized in the progressive randomization, the similarity between these individual networks is lower than the similarity between corresponding surrogates, indicating moderately larger inter-subject variability in structural connectivity than in wiring length distribution. We also observed significant connection specificity in an independent dataset that included 196 healthy subjects aging from 4 to 85 years old. These consistent and robust observations highlight that the connection specificity is a common and fundamental property of healthy brains during different developmental and aging stages.

Network randomization may be considered as the inter-generation alteration driven by environmental and genetic factors in the context of evolution [42]. The resultant landscape paints a portrait of possible evolutionary trajectories of structural connectomes without cost and adaptive constraints. By comparing empirical networks with those generated by progressive randomization, we could obtain insights into adaptively enhancing features of the brain structural network. We observed all individual networks resided within a narrow window with relatively larger intra-group variance in structural connectivity. We speculate that narrow boundaries may reflect the influence of adaptive selection within which network configurations are different due to diverse neurodevelopment, but still conserve critical organizational properties to underpin brain function. This resonates with the finding [26] that network formation is governed by a set of tightly framed parameters in generative network modeling, with subtle variability in these parameters driving neurodevelopmental diversity of macroscopic brain organization. Based on these results, we speculate that the connection specificity may be a heritable organizational feature that contributes to adaptive enhancement. Future work could investigate the heritability of connection specificity and test the relationship between individuals’ cognitive outcomes and subtle differences in specificity degree.

The physical instantiations of structural connectivity inevitably come with the material and metabolic cost [1]. It is widely believed that brain structural networks are organized under a trade-off between the pressure of cost minimization and the requirement of functional complexity [25, 26], which potentially pits the wiring economy against the functional advantages. However, in the process of progressive randomization (Fig. 4), the increment in long-distance connections is accompanied by the gradual disappearance of connection specificity, implying that the relationship between cost constraints and functional demands may not always be competitive. The observation that the wiring length distribution skewed toward short distances promotes the expression of connection specificity raises a possibility that the short-distance preference of structural connectomes may not only be subject to cost constraints but also facilitate the realization of brain function. Furthermore, it is worth exploring which model can better capture the organizational mechanisms of brain structural networks, a trade-off between cost and function, or a balance between diverse function demands.

Although the spatial, topological, and functional architecture of brain structural networks has been fruitfully investigated, it is still challenging to obtain a mechanistic understanding of how these three characteristics are implicated with one another. The connection specificity revealed in this work may be a good breakthrough. It elucidates how brain spatial architecture impinges on interregional communications, associating brain areas’ length profiles with their functionality and incorporating length dispersion and clustering coefficient into a hierarchical structure. Specifically, we found that brain regions’ length profiles were informative indicators of their functional roles, with unimodal regions (especially the somatomotor network) exhibiting low length dispersion and transmodal regions (especially the default mode network) exhibiting high length dispersion. This increasing dispersion of connection length distributions might be motivated by systematic variation in cytoarchitecture [43], intracortical myelination [44], and laminar differentiation [45], supporting the increasingly integrative function from primary sensory regions to transmodal regions. We also found that the clustering coefficient decreased monotonously with increasing length dispersion, mirroring a hierarchical architecture that potentially served for segregated-integrative information processing. Altogether, our work establishes a putative link between brain spatial, topological, and functional characteristics, opening the door to a comprehensive investigation of brain structural networks from different dimensions and aspects. Alterations in connection specificity or changes in structural organizations built on it may be another promising application, with major implications for treating neurological diseases and delaying cognitive deficits associated with aging.

## 4. Methods

### Connectome datasets

We performed all analyses in 2 independent datasets. The main analyses (discovery) were performed in data from the Department of Radiology, University Hospital Center and University of Lausanne (LAU). The age-related analyses were performed in the Nathan Kline Institute (NKI)/Rockland Sample public dataset. In this section, we offered a brief description of each dataset.

#### LAU

The dataset was collected from a cohort of 70 healthy participants (27 females, 28.8±9.1 years old). Informed content approved by the Ethics Committee of Clinical Research of the Faculty of Biology and Medicine, University of Lausanne was obtained from all participants. Diffusion spectrum images (DSI) were acquired on a 3-Tesla MRI scanner (Trio, Siemens Medical, Germany) using a 32-channel head-coil. The protocol was comprised of (1) a magnetization-prepared rapid acquisition gradient echo (MPRAGE) sequence sensitive to white/gray matter contrast (1-mm in-plane resolution, 1.2-mm slice thickness), (2) a DSI sequence (128 diffusion-weighted volumes and a single b0 volume, maximum b-value 8,000 s/mm2, 2.2 × 2.2 × 3.0 mm voxel size), and (3) a gradient echo EPI sequence sensitive to blood oxygen level-dependent (BOLD) contrast (3.3-mm in-plane resolution and slice thickness with a 0.3-mm gap, TR 1,920 ms, resulting in 280 images per participant). Gray matter was divided into 68 brain regions following Desikan-Killiany atlas [46]. These regions were further subdivided into 114, 219, 448, and 1,000 approximately equally sized nodes according to the Lausanne anatomical atlas using the method proposed by [47]. Individual structural networks were constructed using deterministic streamline tractography, initiating 32 streamline propagations per diffusion direction for each white matter voxel [48]. A consensus adjacency matrix that preserved the density and the edge-length distribution of the individual participant matrices [35, 49, 25] was then estimated using a group-consensus approach [49, 50, 51]. For more details regarding network construction see ref [52].

#### NKI

The dataset consisted of 196 participants (aging from 4 to 85 years old). Informed content approved by the Institutional Review Board was obtained from all participants (informed content was also obtained from child participants and their legal guardians). The scan was performed in a Siemens Trio 3T scanner. The protocol consisted of: (1) 10 minute resting state fMRI scan (R-fMRI), (2) 6-direction diffusion tensor imaging (DTI) scan, (3) 64-direction diffusion tensor imaging scan (2mm isotropic), (4) MPRAGE anatomical scan, (5) MPRAGE anatomical scan SHORTER sequence, (6)T2 weighted sequence, (7) A variety of psychiatric, cognitive and behavioral assessments. The structural network composed of 188 ROIs based on the Craddock 200 atlas [53] was obtained by diffusion tensor imaging (DTI).

### Connection similarity

The connectivity profile of a brain area shapes its interaction pattern with other areas and determines its functional contribution in the brain network [54, 55]. Regions with similar connectivity profiles are considered to perform equivalent functional roles, thus contributing similar signals in interregional communication [49, 36, 37, 38]. To measure the relatedness of input and output signals transmitted by different connections of area i, we calculate the cosine similarity of connectivity profiles of neighbors linked by these connections. We define the connectivity profile of area i’s neighbor j as a vector **w_j_** = [**w_j1_**, …, **w_jN_**], where N denotes the network size. The similarity of area i’s neighbors j and k can be estimated as:

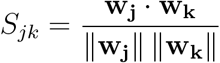

To assess the similarity of different-distance connections, we partition area i’s neighbors into distance bins according to their connection lengths and then constructed a similarity matrix *S^i^* The matrix element 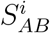 is computed as the average similarity of connectivity profiles of all neighbor pairs (*j, k*) satisfying the condition that j is from the distance bin A and k is from distance bin B. That is, 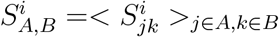. We then repeat the above process for each brain area and average the resultant areal similarity matrices. Finally, we obtain a whole-brain similarity matrix *S* =< *S^i^* >, which measures the extent to which brain areas receive or deliver similar inputs and outputs through structural connections with different distances.

### Connection specificity

We measure the specificity of structural connections as the extent to which different-distance connections lead to neighbors with dissimilar connectivity profiles. Specifically, we construct a matrix *S^d^* to describe the connection length differences across different bins. Let *L_A_* =< *l_A_* > be the average length of connections in bin A, then the element 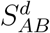 in matrix **S^d^** can be expressed as 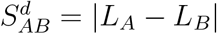. The specificity degree is measured as *r* = –*corr*(*vec*(**S**), *vec*(**S^d^**)), that is, the negative correlation coefficient between the similarity of neighbors’ connectivity profiles and connection length differences. To test whether the specificity of different-distance connections is a nontrivial property of brain networks, we compared the observed similarity matrix and r-value with those of null models. We found that the similarity is significantly negative correlated with the connection length differences, indicating that the connection specificity is a meaningful architecture of human connectomes. Notably, the connection specificity is robust to moderate adjustment of bin number (SI Appendix, Fig. .2).

### Network randomization

We generate four types of null surrogates: (i) Random surrogates(R) that disrupt all network properties except the number of nodes and connections. These surrogates are generated by randomly connecting pairwise brain areas until the connection number equals that of the original network. (ii) Degree-preserving surrogates(D) that restore the degree sequence of empirical network. These surrogates are created using the Brain Connectivity Toolbox https://www.nitrc.org/projects/bct, which is a matlab toolbox for complex network analysis of brain connectivity data. (iii) Geometry-preserving surrogates(G) that restore the wiring length distribution of empirical network. These surrogates are obtained using variants of the randomization scheme in ref [42]. We divide all network connections into a set of equal-width bins by their lengths and then randomly shuffle connections within each bin to ensure the preservation of wiring length distribution in surrogate realizations. In our datasets, 50 bins are sufficient for fine-grained decomposition of connection lengths. (iv) Degree-geometry surrogates (DG) that simultaneously restore original degree sequence and wiring length distribution. The algorithm of generating these surrogates is similar to that of degree-preserving surrogates, with the important exception that we only break and rewire connections within the same bins. When introducing randomization progressively, we only randomly selected a subset of network connections to be shuffled.

### Length dispersion

All interregional connections in the brain network are divided into M bins by their lengths. For any brain area i, let 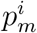 be the probability that it possesses connections in bin *m*. Then define the length dispersion of area i as:

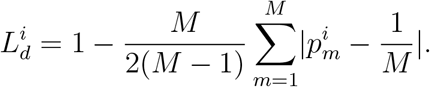

That is, the similarity between area i’s connection length distribution and the uniform distribution. The 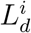 is bounded in [0,1]. At the extreme left of the interval, where 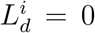, all connections of area i fall in one distance bin. These connections with roughly equivalent lengths may confer area i highly specialized function by transmitting a set of similar inputs and outputs. At the opposite extreme, where 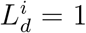, area i possessed connections that were distributed equally among all distance bins. These connections with different lengths render area i the capacity to exert influence to and receive influence from neighbors with dissimilar connectivity profiles, providing an anatomical substrate for the integration of distributed and diverse information. Therefore, the length dispersion may be an informative indicator of the interregional communication patterns, mirroring the functional roles of regions within the brain network.

### Network hierarchy

We analyzed the hierarchical structure of brain networks following the basic idea in ref [56, 57, 58]. In our work, the hierarchy measures the tendency that nodes with high length dispersion exhibit low clustering coefficients. The hierarchy index *β*_1_ is estimated by fitting the relationship 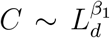, where *C* is the clustering coefficient. We can also simplify the hierarchy index as the negative correlation coefficient between length dispersion and clustering coefficient: 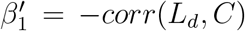. This simplified index is strongly correlated to the index *β*_1_ (SI Appendix, Fig. .3).

## Supplementary information

The robustness analysis are provided in Supplementary Materials.

## Acknowledgments

This work is supported by Program of National Natural Science Foundation of China Grant No. 42050105, 11871004, 11922102, 62141605 and National Key Research and Development Program of China Grant No. 2018AAA0101100

## Declarations

Not applicable

**SI Appendix, Fig .1:**
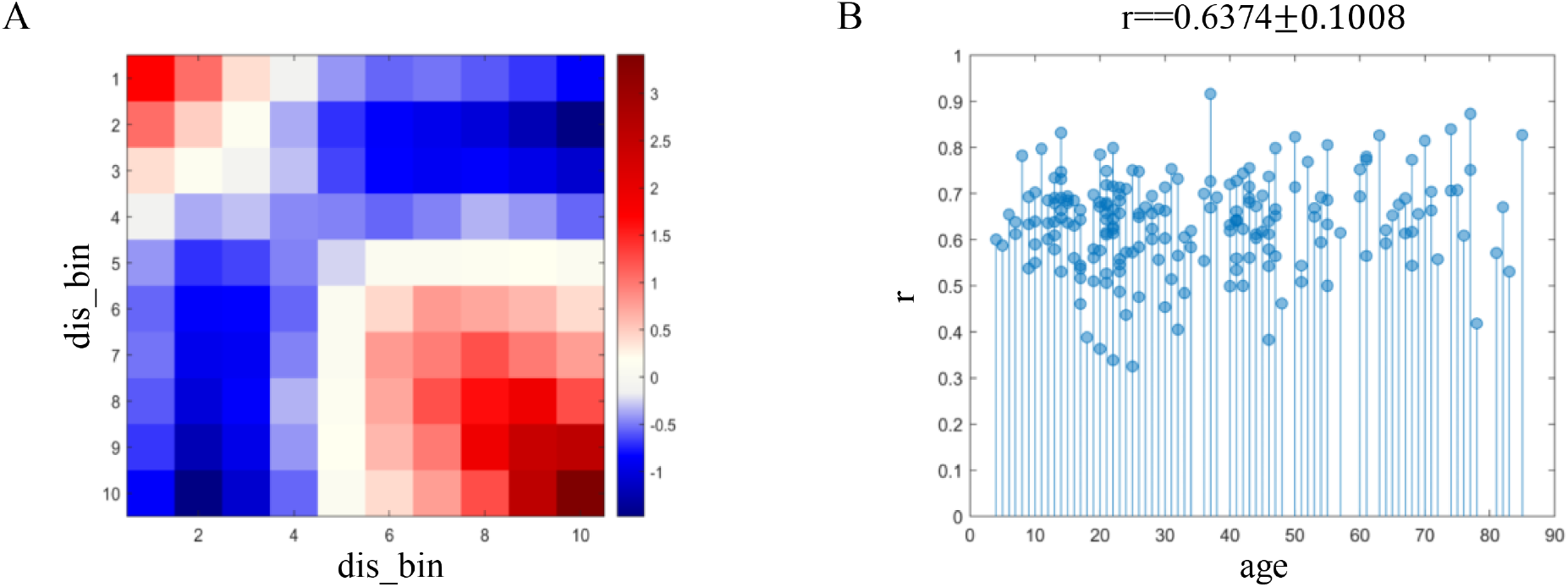
Connection specificity in an independent dataset. A. The average similarity matrix in the Nathan Kline Institute (NKI)/Rockland Sample public dataset. B. The specificity of 196 individuals in the NKI dataset.

**SI Appendix, Fig .2:**
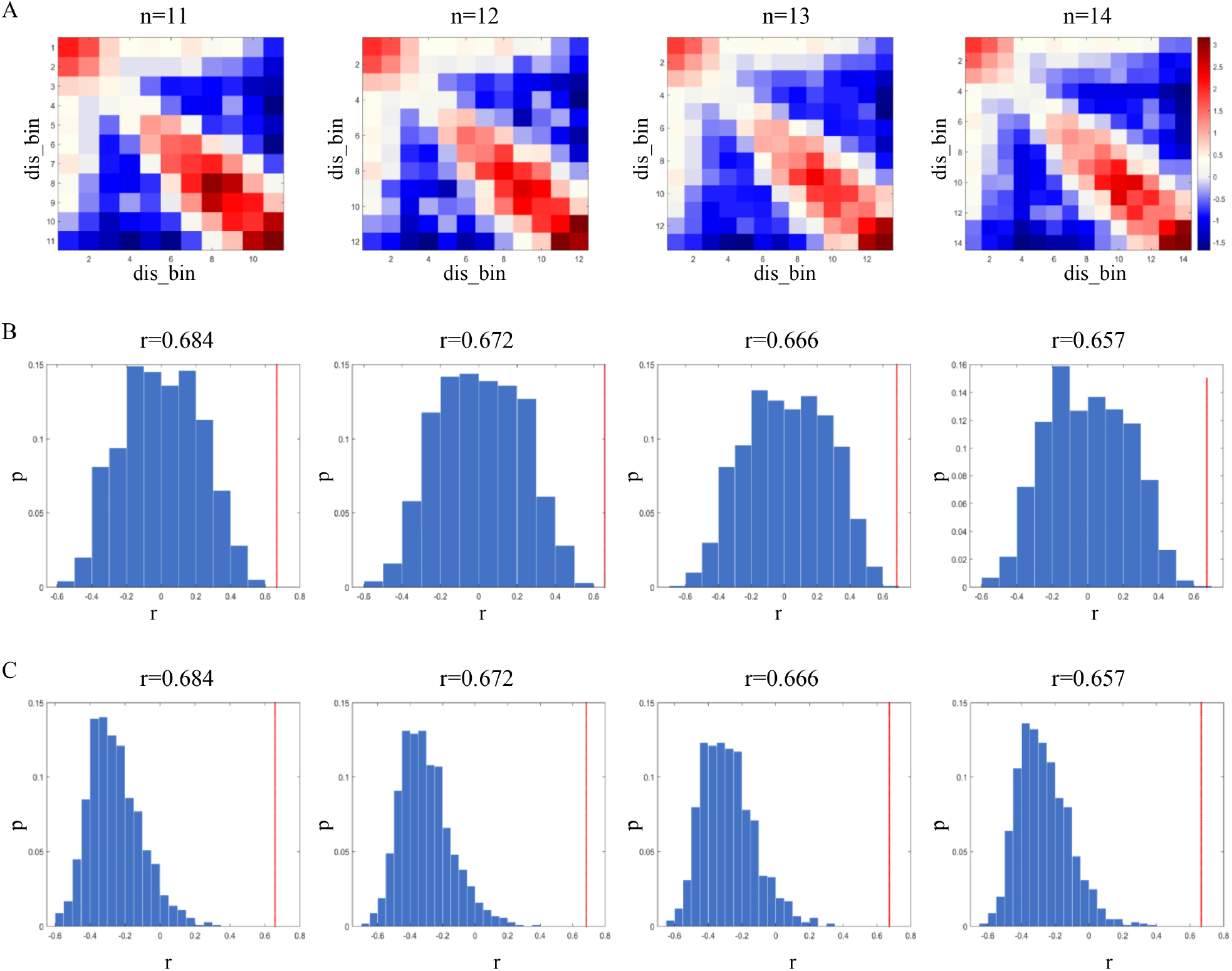
Robustness of bin number. A. The similarity matrix with different numbers of distance bins. B. Comparison between empirical values with the null distribution generated by randomly permuting areas’ spatial locations while preserving network topology. C. Comparison between empirical values with the null distribution generated by randomly rewiring network edges while preserving degree sequence.

**SI Appendix, Fig .3:**
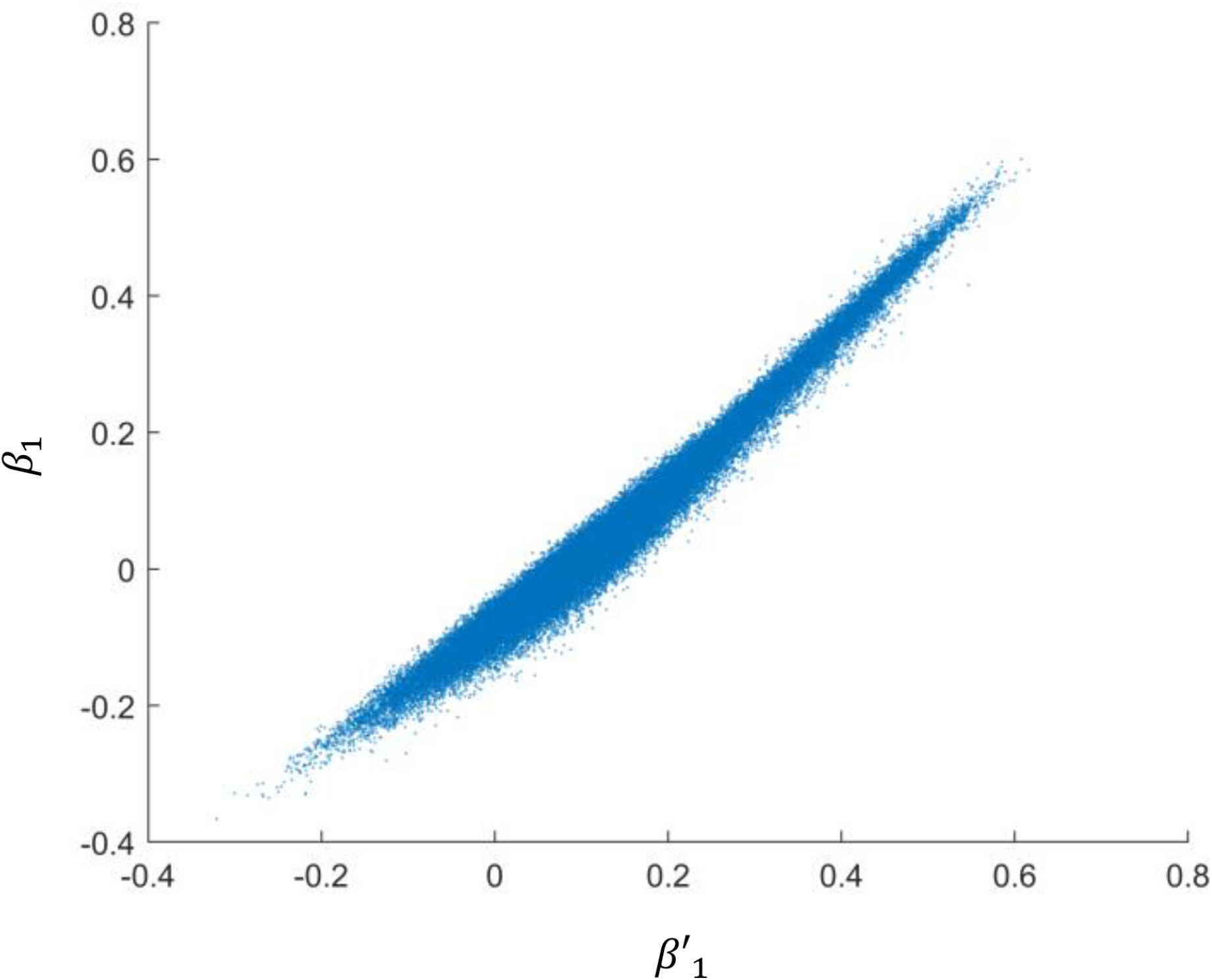
: Correlation between hierarchy index *β*_1_ and 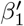. We computed *β*_1_ and 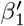 of 100,000 randomized surrogates and found a strong correlation between these two hierarchy indices

